# Simultaneous Brain, Brainstem and Spinal Cord pharmacological-fMRI reveals endogenous opioid network interactions mediating attentional analgesia

**DOI:** 10.1101/2021.05.05.442823

**Authors:** Valeria Oliva, Ron Hartley-Davies, Rosalyn Moran, Anthony E. Pickering, Jonathan C.W. Brooks

**Affiliations:** School of Physiology, Pharmacology & Neuroscience, Biomedical Sciences Building, University of Bristol, Bristol, BS8 1TD, UK; Clinical Research and Imaging Centre, School of Psychological Science, University of Bristol, Bristol, BS8 1TU, UK; Medical Physics, University Hospitals Bristol & Weston NHS Trust, BS2 8HW, UK; Department of Neuroimaging, Institute of Psychiatry, Psychology & Neuroscience, King’s College London, SE5 8AF, UK; Anaesthesia, Pain & Critical Care Sciences, Bristol Medical School, University Hospitals Bristol & Weston, Bristol, BS2 8HW, UK; University of East Anglia Wellcome Wolfson Brain Imaging Centre, School of Psychology, Lawrence Stenhouse Building, Chancellors Drive, NR4 7TJ, UK; Department of Anesthesiology, University of California San Diego, 9500 Gilman Dr, La Jolla, CA 92093, USA

## Abstract

Pain perception is decreased by shifting attentional focus away from a threatening event. This attentional analgesia engages parallel descending control pathways from anterior cingulate (ACC) to locus coeruleus, and ACC to periaqueductal grey (PAG) – rostral ventromedial medulla (RVM), indicating possible roles for noradrenergic or opioidergic neuromodulators. To determine which pathway modulates nociceptive activity in humans we used simultaneous whole brain-spinal cord pharmacological-fMRI (N=39) across three sessions. Noxious thermal forearm stimulation generated somatotopic-activation of dorsal horn (DH, C6 segment) whose activity mirrored attentional pain modulation. Activity in an adjacent cluster reported the interaction between task and noxious stimulus. Effective connectivity analysis revealed that ACC recruits PAG and RVM to modulate spinal cord activity. Blocking endogenous opioids with Naltrexone impairs attentional analgesia and disrupts RVM-DH and ACC-PAG connectivity. Noradrenergic augmentation with Reboxetine did not alter attentional analgesia. Cognitive pain modulation is mediated by opioidergic ACC-PAG-RVM descending control which supresses spinal nociceptive activity.

## Introduction

Pain is a fundamental and evolutionarily-conserved cognitive construct that is behaviourally prioritised by organisms to protect themselves from harm and facilitate survival. As such pain perception is sensitive to the context within which potential harm occurs. There are well recognised top-down influences on pain that can either suppress (e.g. placebo (*1*) or task engagement (*2*)) or amplify (e.g. catastrophising (*3*), hypervigilance (*4*) or nocebo (*5*)) its expression. These processes influence both acute and chronic pain and provide a dynamic, moment by moment regulation of pain as an organism moves through their environment.

A simple shift in attention away from a noxious stimulus can cause a decrease in pain perception – a phenomenon known as attentional analgesia. This effect can be considered to be a mechanism to enable focus, allowing prioritisation of task performance over pain interruption (*6, 7*). This phenomenon is reliably demonstrable in a laboratory setting (*8*) and a network of cortical and brainstem structures have been implicated in attentional analgesia (*9–17*).

We have shown that two parallel pathways are implicated in driving brainstem activity related to attentional analgesia (*16, 18*). Projections from rostral anterior cingulate cortex (rACC) were found to drive the periaqueductal grey (PAG) and rostral ventromedial medulla (RVM), which animal studies have shown to work in concert using opioidergic mechanisms to regulate spinal nociception (*19–22*). Similarly, a bidirectional connection between rACC and locus coeruleus (LC) was also directly involved in attentional analgesia. As the primary source of cortical noradrenaline, the LC is thought to signal salience of incoming sensory information (*23, 24*), but can also independently modulate spinal nociception (*25–28*). Although these animal studies provide a framework for our understanding of descending control mechanisms that are likely to be mediating attentional analgesia, the network interactions between brain, brainstem and spinal cord and the neurotransmitter systems involved in producing attentional analgesia have yet to be elucidated in humans. In part, this gap in our knowledge is because of the distributed extent of the network spanning the entire neuraxis from forebrain to spinal cord, which has not been amenable to simultaneous imaging approaches in humans.

To address this issue, we conducted a double-blind, placebo-controlled, three arm, cross-over pharmacological-fMRI experiment to investigate attentional analgesia using whole neuraxis imaging and a well validated experimental paradigm. To engage attention, we utilised a rapid serial visual presentation (RSVP) task (*16, 18, 29*) with individually calibrated task difficulties (easy or hard), which was delivered concurrently with thermal stimulation adjusted per subject to evoke different levels of pain (low or high). We took advantage of recent improvements in signal detection (*30*) and pulse sequence design to simultaneously capture activity across the brain, brainstem, and spinal cord (i.e. whole central nervous system, CNS) in a single contiguous functional acquisition with slice-specific z-shimming (*31–33*). To resolve the relative contributions from the opioidergic and noradrenergic systems, subjects received either the opioid antagonist naltrexone, the noradrenaline re-uptake inhibitor reboxetine, or placebo. By measuring the influence of these drugs on pain perception, BOLD activity and effective connectivity between a priori specified regions known to be involved in attentional analgesia (rACC, PAG, LC, RVM, spinal cord (*16–18*)), we sought to identify the network interactions and neurotransmitter mechanisms mediating this cognitive modulation of pain.

## Results

A total of 39 subjects (mean age 23.7, range [18 - 45] years, 18 females) completed the fMRI imaging sessions with a 2*2 factorial experimental design (RSVP task easy / hard and high / low thermal stimulus) with a different drug administered orally before each session (naltrexone (50mg), reboxetine (4mg) or placebo). The behavioural signature of attentional analgesia is a task*temperature interaction, driven by a reduction in pain ratings during the high temperature-hard task condition (*16, 18, 34*). A first level analysis of the pooled pain data across all experimental sessions showed: a main effect of temperature (F (1,38) = 221, P=0.0001) with higher scores under the high temperature condition; a main effect of task (F (1,38) = 4.9, P=0.03); and importantly demonstrated the expected task*temperature interaction consistent with attentional pain modulation (F (1, 38) = 10.5, P = 0.0025, Supplementary Figure 1).

To assess the impact of the drugs on attentional analgesia, each experimental session was analysed independently (Figure 1A). Attentional analgesia was seen in the placebo condition (task*temperature interaction (F (1, 38) = 11.20, P = 0.0019), driven primarily by lower pain scores in the hard|high vs easy|high condition (37.5±19.4 vs 40.4±19.8, mean±SD, P = 0.001). Similarly, subjects given Reboxetine showed a task*temperature interaction (F (1, 38) = 9.023, P = 0.0047), again driven by decreased pain scores in the hard|high vs easy|high condition (31.9 ± 15.8 vs 35.6 ± 15.5, P = 0.0034). In contrast, Naltrexone blocked the analgesic effect of attention with no task*temperature interaction (F (1, 38) = 0.4355, P = 0.5133, hard|high (37.4±17.1) vs easy|high (38.3±17.1)). This effect was specific to attentional analgesia as neither drug had any effect on the calibrated temperature for the high thermal stimulus or the speed of character presentation for the RSVP task (Supplementary Figure 2). Behaviourally these findings indicate that the attentional analgesic effect is robust, reproducible between and across subjects and that it involves an opioidergic mechanism.

**Figure 1.**
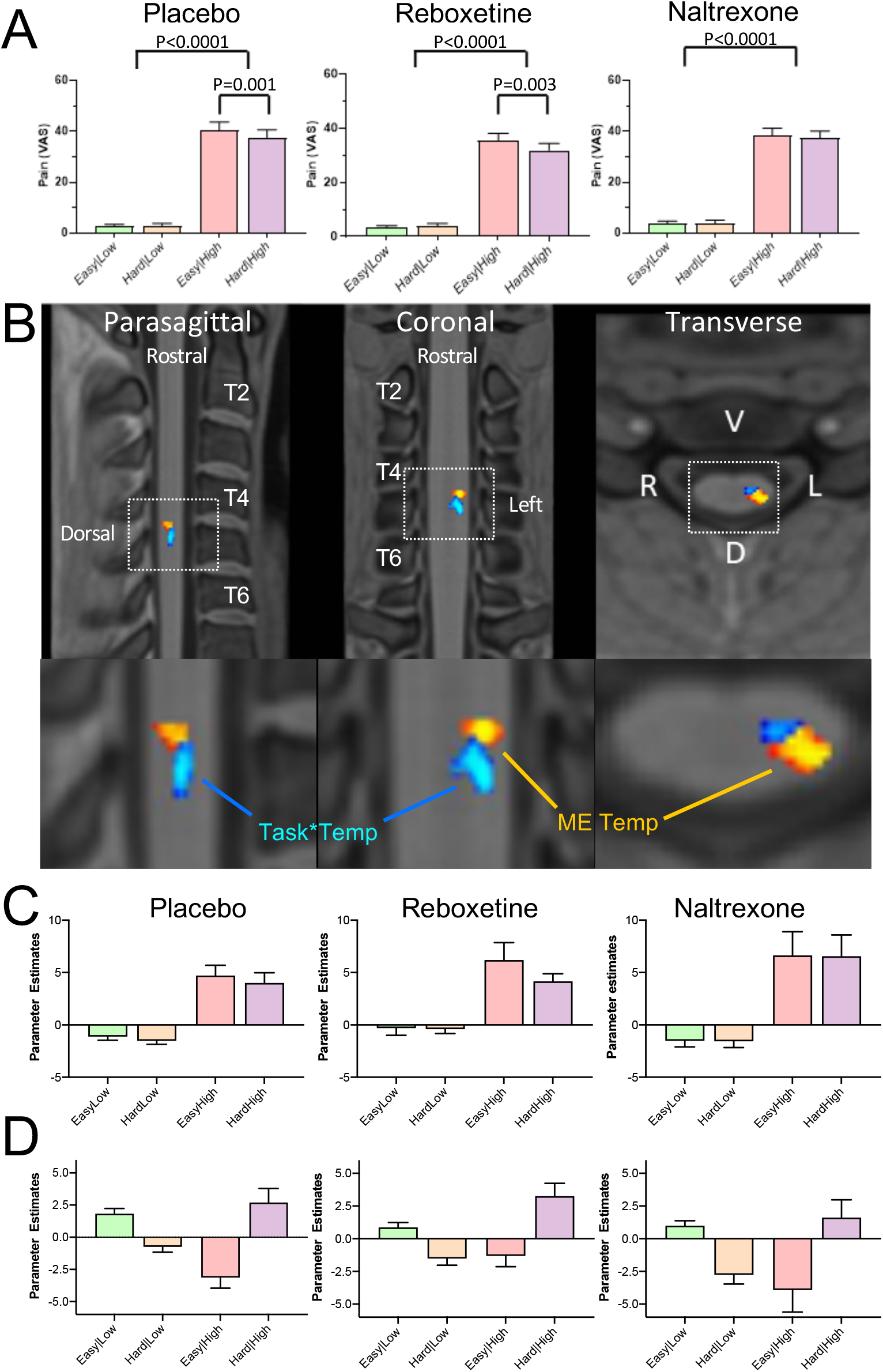
Main effect of temperature and task*temperature interaction in the spinal cord. (A) Pain scores across the four experimental conditions (i.e. easy|low, hard|low, easy|high and hard|high), for the three drugs. All conditions showed a main effect of temperature (Two-way mixed effects ANOVA). Attentional analgesia was seen in the placebo and reboxetine limbs with a task*temperature interaction (F (1, 38) = 11.20, P = 0.0019 and F (1, 38) = 9.023, P = 0.004 respectively). In both cases this was driven by lower pain scores in the hard|high versus easy|high condition (Sidak’s post hoc test). In contrast Naltrexone blocked the analgesic effect of attention as reflected in a loss of the task*temperature interaction (F (1, 38) = 0.4355, P = 0.5133). (B) Cervical spine fMRI revealed two distinct clusters of activity within the left side of the C6 cord segment. The first showing the main effect of temperature (red-yellow, *Spinal_noci_*) and a second showing task*temperature interaction (blue-light blue, *Spinal_int_*) (significance reported with P<0.05 (TFCE) within a left sided C5/C6 anatomical mask). No cluster reached significance for the main effect of task. (C) Parameter estimates from the *Spinal_noci_* cluster revealed a decrease in BOLD in the hard|high versus easy|high condition, seen in placebo and reboxetine arms but not in naltrexone. Note the similarity in pattern with the pain scores in (A). (D) Extraction of parameter estimates from the *Spinal_int_* cluster revealed an increase in BOLD in the hard|high condition, across all three drug sessions but this was attenuated in the Naltrexone condition. Mean +SEM. Parameter estimates extracted from the peak voxel in each cluster.

We also noted a drug*temperature interaction on pain ratings in the first level analysis (F (2, 76) = 3.2, P = 0.04, Supplementary Figure 1). Comparing reboxetine versus placebo showed the presence of a drug*temperature interaction (F (1, 38) = 5.060, P = 0.03), with lower pain scores in the presence of reboxetine indicating that it was underpinned by an analgesic effect of the noradrenergic reuptake inhibitor (in contrast naltrexone vs placebo showed no drug*temperature interaction).

### Whole CNS fMRI: main effects and interactions

To determine the neural substrates for attentional analgesia and to identify the possible involvement of the noradrenergic and opioidergic systems, we initially defined a search volume in which to focus subsequent detailed fMRI analyses. This was achieved by pooling individually averaged data across the three experimental imaging sessions to estimate main effects and interactions across all levels of the neuraxis.

#### Spinal cord

A cluster of activation representing the positive main effect of temperature was identified in the left dorsal horn (DH), in the C6 spinal segment (Figure 1B). This represents activity in a population of neurons that responded more strongly to noxious thermal stimulation. This *Spinal_noci_* cluster was somatotopically localised, given that the thermal stimuli were applied to the left forearm in the C6 dermatome. BOLD parameter estimates were extracted to investigate the activity of this *Spinal_noci_* cluster across the four experimental conditions and three drug sessions (Figure 1C). In the placebo session, the pattern of BOLD signal change across conditions was strikingly similar to the pain scores (Figure 1A), and the response to a noxious stimulus was lower in the hard|high than easy|high condition, indicating that the *Spinal_noci_* activity was modulated during attentional analgesia. The same pattern was evident in the reboxetine condition, again indicating spinal cord modulation during attentional analgesia. However, this was not observed in the naltrexone arm, where the *Spinal_noci_* cluster showed a similar BOLD response in the easy|high and hard|high conditions, again mirroring the pain ratings and consistent with the opioid antagonist-mediated blockade of attentional analgesia.

Analysis of task*temperature interaction revealed a second discrete spinal cluster (Spinal_int_, Figure 1B). This was also located on the left side of the C6 segment but was slightly caudal, deeper and closer to the midline with respect to the *Spinal_noci_* cluster (with only marginal overlap). Extraction of BOLD parameter estimates from the *Spinal_int_* cluster in the placebo and reboxetine condition, showed an increased level of activity in the hard|high condition (Figure 1D). However, this was not evident in the presence of naltrexone. This activity profile suggests this *Spinal_int_* cluster, potentially composed of spinal interneurons, plays a role in the modulation of nociception during the attentional analgesic effect.

#### Brainstem and whole brain

To identify the regions of the brainstem involved in mediating attentional analgesia and potentially interacting with the spinal cord, a similar pooled analysis strategy was employed. Analysis of the main effect of temperature within a whole brainstem mask showed substantial clusters of activity in the midbrain (PAG) and medulla (RVM) with more discrete clusters in the dorsal pons bilaterally (LC) (Figure 2A). In the main effect of task, the pattern of brainstem activation was more diffuse (Figure 2A), but again included activation of the PAG, RVM and bilateral LC. Importantly for the mediation of attentional analgesia, no task*temperature interaction was observed within the brainstem.

**Figure 2.**
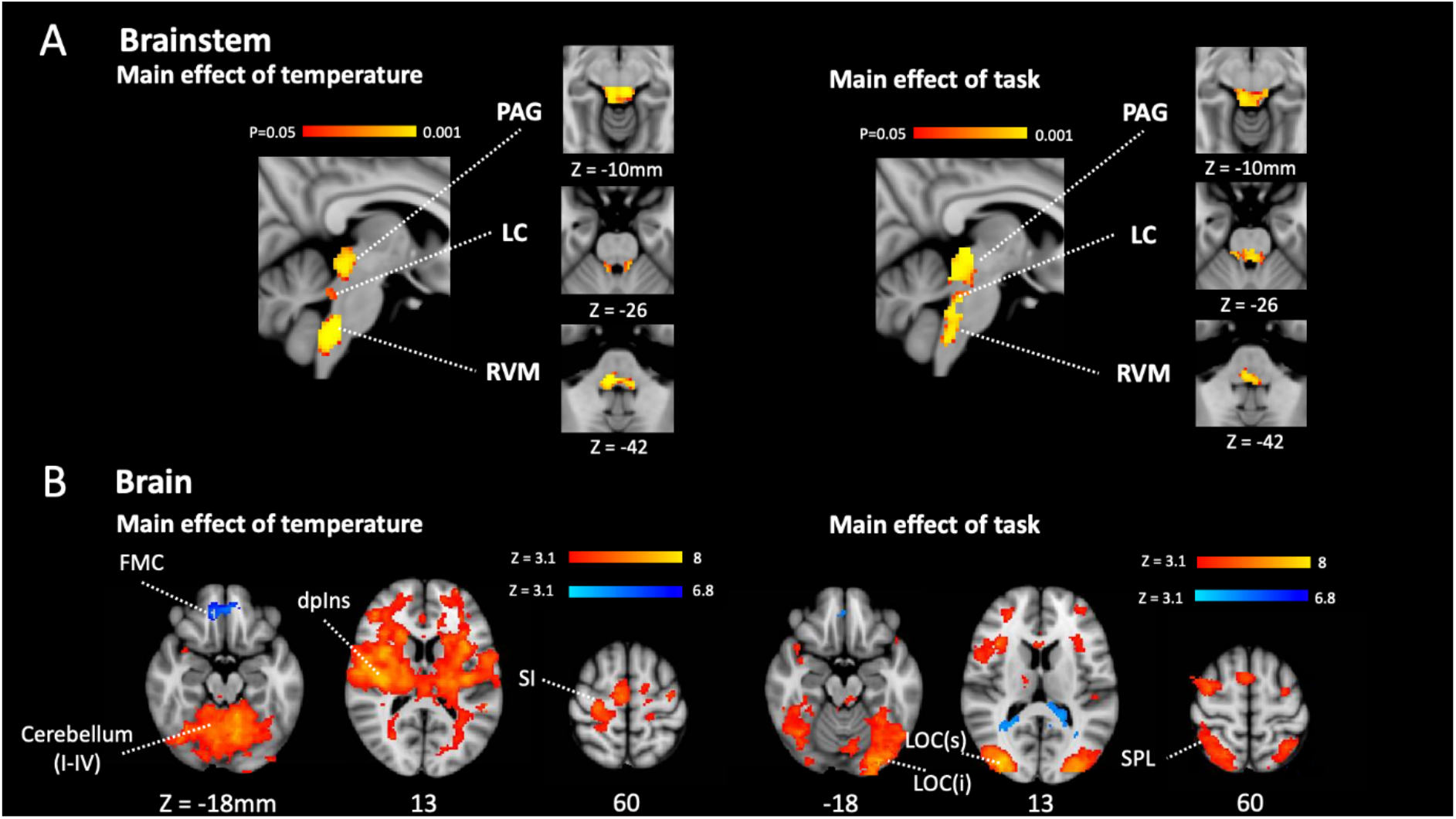
Main effect of task and temperature in Brainstem and Cerebrum. (A) Main effect of temperature and task in the brainstem after permutation testing with a whole brainstem mask showing clusters of activation in PAG, bilateral LC and RVM. Activity reported for P<0.05 (TFCE).(B) Main effects of temperature and task in brain. In the main effect of temperature contrast there were clusters of activation in a number of pain related sites including in the contralateral primary somatosensory cortex, the dorsal posterior insula and the PAG (red-yellow). The frontal medial cortex de-activated (blue-light blue). In the main effect of task contrast there were clusters of activation in the visual and attention networks including superior parietal cortex, the frontal pole, and the anterior cingulate cortex (red-yellow). The posterior cingulate cortex and lateral occipital cortex showed de-activation (blue-light blue). Activity was estimated with a cluster forming threshold of Z>3.1 and corrected significance level of P<0.05.

At the brain level, analysis of the main effect of temperature contrast showed activation in pain-associated regions such as primary somatosensory cortex, dorsal posterior insula, operculum, anterior cingulate cortex and cerebellum with larger clusters contralateral to the side of stimulation (i.e. right side of brain). A cluster in the medial pre-frontal cortex exhibited deactivation. (Figure 2B, Supplementary Table 1). For the main effect of task, bilateral activation was seen in attention and visual processing areas including lateral occipital cortex, anterior insular cortex and anterior cingulate cortex. Deactivation was observed in the cerebellum (Crus I), precuneus and lateral occipital cortex (superior division). (Figure 2B, Supplementary Table 1). No cluster in the whole brain analysis reached significance in the positive task*temperature interaction.

The distribution of these patterns of regional brain and brainstem activity were closely similar to those found in our previous studies of attentional analgesia (*16, 18, 34*) but with the difference that no area in the brain or brainstem showed a task*temperature interaction (unlike the spinal cord). This motivated a network connectivity analysis (*18*) to determine which regions were communicating the effect of the cognitive attentional task to the spinal cord.

(PAG – Periaqueductal grey, LC – Locus coeruleus, RVM – Rostral ventromedial medulla, FMC – Frontomedial cortex, dpIns – dorsal posterior insula, S1 – primary somatosensory cortex, LOC – Lateral ocipital cortex (sup and inf), SPL Superior parietal lobule.)

### Attentional analgesia and effective network connectivity

To define an attentional analgesia network, we performed a generalised psychophysiological interaction (gPPI) analysis for the placebo condition alone within the *a priori* identified seed/target regions (after (*18*)): ACC, PAG, right LC and RVM to which we added the cervical spinal cord (left C5/C6 mask).

The gPPI identified the following pairs of connections [<u>seed</u>-target] as being significantly modulated by our experimental conditions (Figure 4A):

- main effect of temperature [<u>PAG</u>-rLC], [<u>rLC</u>-ACC], [<u>rLC</u>-RVM] and [<u>RVM</u>-DH]
- main effect of task [<u>RVM</u>-rLC] and [<u>PAG</u>-ACC]
- task*temperature interaction [<u>RVM</u>-PAG], [<u>RVM</u>-rLC], and [<u>RVM</u>-DH].

This pattern of network interactions has a number of common features shared with our previous analysis (*18*) including the task modulation of connectivity between PAG and ACC and the effect of the task*temperature interaction on connectivity between RVM and PAG. The new features were the influence of all conditions on communication between RVM and rLC and the important linkage between the dorsal horn (DH) activity and RVM which is modulated by both temperature and the task*temperature interaction.

Parameter estimates extracted from the connections modulated by task, revealed that the <u>PAG</u>-ACC, <u>RVM</u>-PAG, <u>RVM</u>-rLC, and <u>RVM</u>-DH connections were stronger in the hard|high versus the easy|high condition, consistent with their potential roles in attentional analgesia (Figure 3).

**Figure 3.**
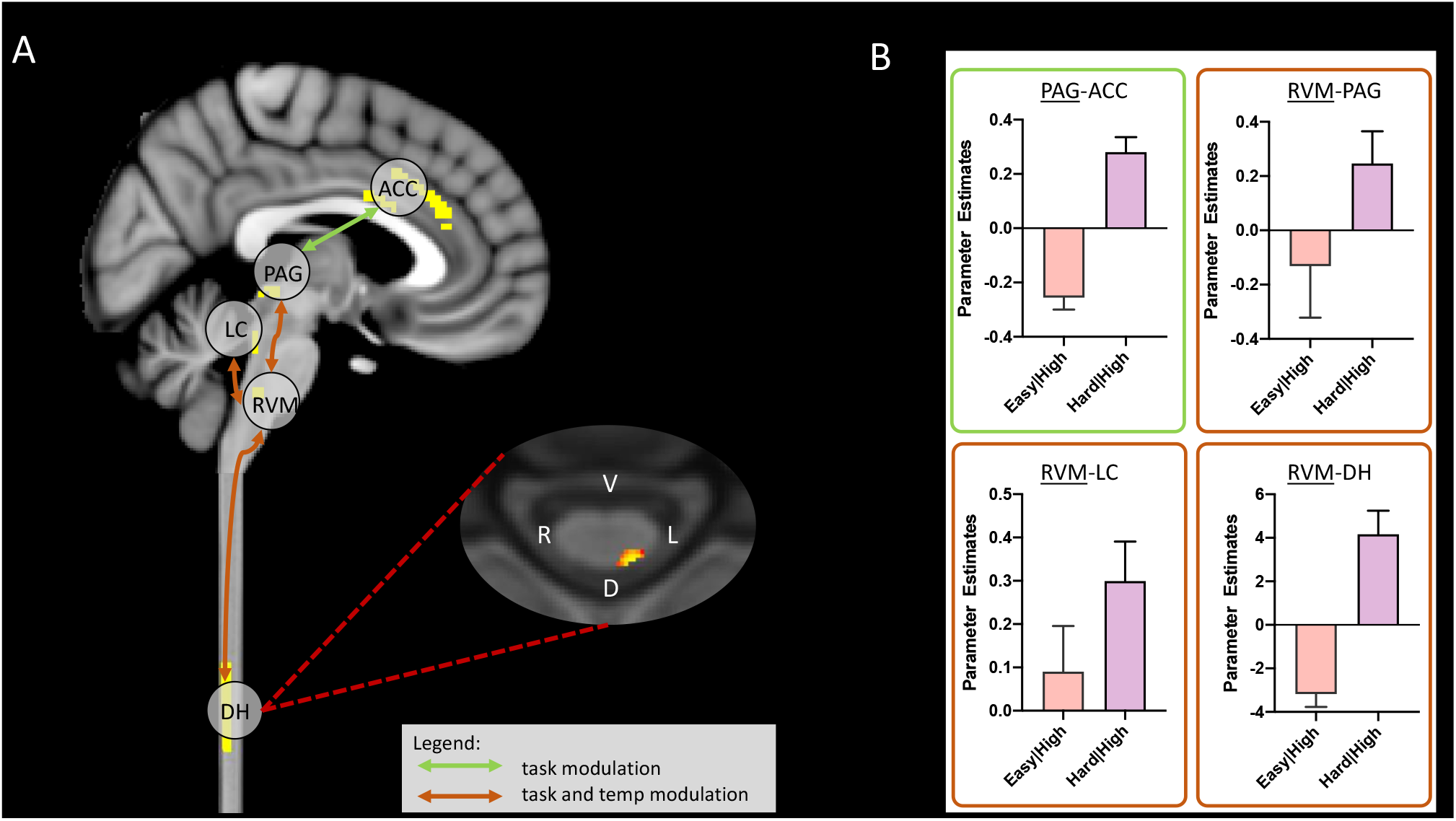
Summary of significant connection changes revealed by the gPPI analysis that were modulated by task (placebo condition only). (A) Permutation testing revealed a significant change in connectivity in the main effect of task contrast between ACC and PAG, and in the task*temperature interaction contrast between PAG and RVM, LC and RVM, and importantly RVM and DH. Masks used for time-series extraction are shown in the sagittal slice (yellow). The spinal cord axial slice shows the voxels with significantly connections with RVM (threshold at P = 0.1 for visualization purposes). (B) Extraction of parameter estimates revealed an increase in coupling in the analgesic condition for all of these connections (i.e. hard|high). (Mean ± SEM).

### Impact of neuromodulators on regional brain activations and network interactions

Having identified this group of regions, in a network spanning the length of the neuraxis, whose activity and connectivity correspond to aspects of the attentional analgesia paradigm we examined whether naltrexone or reboxetine affected the regional BOLD activity or connectivity, comparing each drug against the placebo condition (using paired t-tests).

At the whole brain level, neither drug altered the activations seen for the main effect of temperature. Only the left anterior insula responded more strongly in the presence of Naltrexone for the main effect of task (Supplementary Figure 3B), however this was not considered relevant to the analgesic effect as our behavioural findings showed no effect of naltrexone on task performance (Supplementary Figure 2B). In the brainstem, a stronger response to temperature was detected in the lower medulla in the presence of naltrexone compared to placebo (Supplementary Figure 3B). There was no difference between naltrexone and placebo in the main effect of task in the brainstem. Similarly, no differences in either main effect were uncovered in the brainstem for the reboxetine versus placebo comparison.

The relative lack of effect of either drug on the net changes in regional BOLD provided little evidence for the localisation of their effects in either blocking attentional analgesia (naltrexone) or producing antinociception (reboxetine). However, it has previously been demonstrated that administration of opioidergic antagonists such as naloxone have measurable effects on neural dynamics assessed with fMRI (e.g. (*35*)). Therefore, we investigated the network of brain, brainstem and spinal regions that show effective connectivity changes associated with attentional analgesia and explored whether these patterns were altered in the presence of reboxetine or naltrexone (paired t-tests versus placebo).

The administration of naltrexone, which abolished attentional analgesia behaviourally, significantly reduced the connection strength of <u>RVM</u>-DH in the task*temperature interaction (Figure 4), indicating a role for opioids in this network interaction. The communication between ACC and PAG was also significantly weakened by both naltrexone and reboxetine, suggesting this connection to be modulated by both endogenous opioids and noradrenaline (Figure 4). The strength of the <u>RVM</u>-LC connection in the main effect of temperature was significantly diminished by reboxetine. None of the other connections in the network were altered significantly by the drugs compared to placebo.

**Figure 4.**
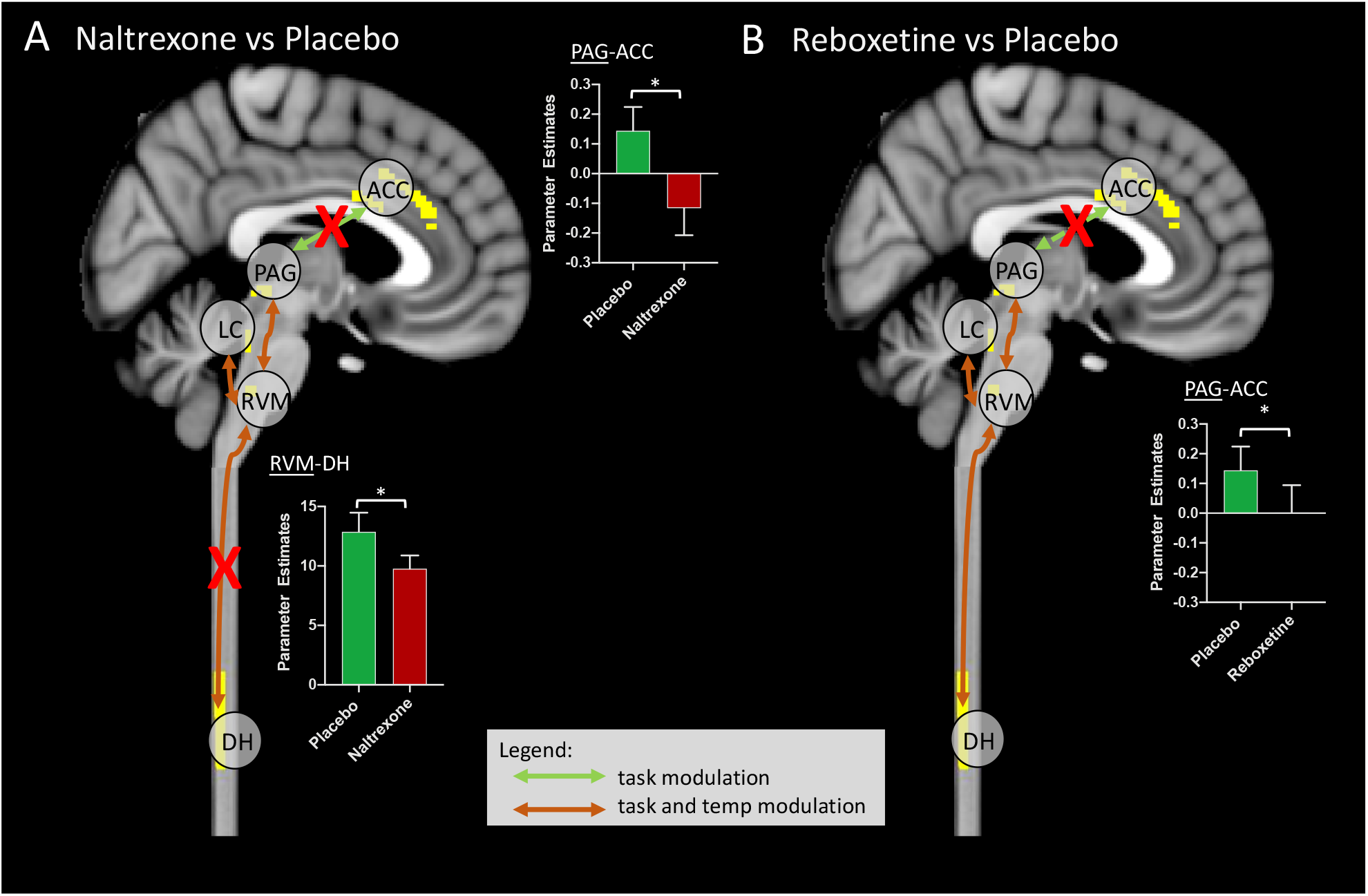
Alteration of functional connectivity after dosing with naltrexone or reboxetine compared to placebo. The ACC-PAG connection was significantly weakened by Naltrexone and Reboxetine administration. The RVM-DH connection was significantly weakened by Naltrexone. Red crosses indicate significantly weaker connections after drug. Inset bar plots show BOLD parameter estimates extracted from the <u>PAG</u>-ACC and <u>RVM</u>-DH connections. (Means±SEM, paired t-test, *P<0.05).

## Discussion

Using brain, brainstem and spinal cord fMRI we have been able to simultaneously measure the changes in neural activity during this attentional pain modulation study at all levels of the neuraxis during a randomised, placebo-controlled, crossover pharmacological study. This approach allowed unambiguous identification of the nociceptive signal at its site of entry in the dorsal horn and revealed that the task-driven cognitive reductions in pain perception echo the change in absolute BOLD signal at a spinal level. Remarkably the spinal imaging also identified a nearby cluster of neural activity that tracked the interaction between cognitive task and thermal stimulus. Analysis of effective connectivity between brain and brainstem regions and the spinal cord in a single acquisition allowed extension from previous findings (*16–18, 34, 36*) to demonstrate causal changes mediating the interaction of pain and cognitive task including descending influences on the spinal dorsal horn. Naltrexone selectively blocked attentional analgesia through reduced connectivity between RVM and dorsal horn and well as between ACC and PAG. This provides a compelling demonstration of the opioid-dependent mechanisms in the descending pain modulatory pathway that is recruited to mediate the attentional modulation of pain.

The use of individually titrated noxious and innocuous stimuli from a thermode applied to the C6 dermatome of the medial forearm, allowed the identification of a somatotopic *Spinal_noci_* cluster in the main effect of temperature contrast in the dorsal horn of the C6 segment. This was strikingly similar to the pattern of activation noted in several previous focussed spinal imaging pain studies in humans (*17, 36–38*) and non-human primates (*39*) which gives additional confidence in its identification as reflecting the nociceptive input. The pattern of extracted absolute BOLD from the *Spinal_noci_* cluster across the four experimental conditions closely paralleled the changes in pain percept as it was modulated by task. This is similar to the seminal findings from electrophysiological recordings in non-human primates (*40*), which showed thermal stimulus evoked neural activity in the spinal nucleus of the trigeminal nerve to be altered by attentional focus. Further, it suggested that task related modulation of pain (*8*) could occur at the first relay point in the nociceptive transmission pathway. This finding of cognitive modulation of nociceptive input was extended through human spinal fMRI by Sprenger and colleagues (*17*), who in a second psychophysical experiment with naloxone provided evidence that the modulation of pain percept may involve opioids. We show that naltrexone attenuates spinal responses to attentional analgesia, which underly the behavioural differences between the high|hard and easy|hard conditions.

Uniquely, our 2×2 factorial study design enabled the identification of neural activity reading out the interaction between task and temperature which strikingly was only seen at a spinal level in a cluster located deep and medial to the *Spinal_noci_* cluster. The activity in this *Spinal_int_* cluster was highest in the high|hard condition. This would be consistent with the presence of a local interneuron population in the deeper dorsal horn that could influence the onward transmission of nociceptive information (*41, 42*). Such a circuit organisation is predicted by many animal models of pain regulation with the involvement of inhibitory interneurons that shape the incoming signals from the original gate theory of Melzack and Wall (*43*) through to descending control (*44*). For example, opioids like enkephalin are released from such local spinal inter-neuronal circuits (*45, 46*) and similarly descending noradrenergic projections exert their influence in part via inhibitory interneurons and an alpha1-adrenoceptor mechanism (*47–50*). As such the ability to resolve this *Spinal_int_* cluster opens a window into how such local interneuron pools may be recruited to shape nociceptive transmission in humans according to cognitive context.

Since our goal was to explore the functional connections between brain, brainstem, and spinal cord, we opted to use a single acquisition, with identical imaging parameters (e.g. orientation of slices, voxel dimensions, point spread function) for the entire CNS. This differs from other approaches (*31–33, 36*), and is motivated by the idea that the use of different acquisition parameters for brain and spinal cord could be a confounding factor in connectivity analyses. By taking advantage the z-shimming approach (*31*) and of the recently developed Spinal Cord Toolbox (*51*), we have been able to detect significant BOLD signal changes in response to experimental manipulations. A key objective of the study was to determine how the information regarding the attentional task demand could be conveyed to the spinal cord. Analysis of regional BOLD signal showed activity in both the main effect of task and of temperature in all three of the key brainstem sites PAG, RVM and LC with no interaction between task and temperature in the brainstem providing little indication as to which area might be mediating any analgesic effect (in line with previous (*18*)). However, an interaction effect was observed on the effective connectivity between RVM and dorsal horn, with coupling highest in high|hard conditions. The importance of this descending connection to the attentional analgesic effect is emphasised by effect of naltrexone which blocked both the modulation of RVM-DH connectivity and attentional analgesia (a behavioural finding previously noted by Sprenger et al (*17*)). This fits with the classic model of descending pain modulation that has been developed through decades of animal research (*19, 22*) that is engaged in situations of fight or flight and also during appetitive behaviours like feeding and reproduction. However, here we identify that the opioidergic system is also engaged moment by moment in specific contexts during a relatively simple cognitive tasks and uncover one of its loci of action in humans.

Analysis of effective connectivity also showed evidence for modulation of pathways from ACC to PAG and PAG to RVM by task and the interaction between task and temperature, respectively (in agreement with (*18*)). The communication between ACC and PAG was also disrupted by the opioid antagonist naltrexone. This is similar to the previous finding from studies of placebo analgesia where naloxone was shown to disrupt ACC-PAG communication which was also linked to the mediation of its analgesic effects (*35*). Activation of an analogous ACC-PAG pathway in rats has recently been shown to produce an analgesic effect mediated via an inhibition of activity at a spinal level indicating that it indeed represents a component of the descending analgesic system (*52*). Interestingly this study also found that this system failed in a chronic neuropathic pain model. This provides evidence for top-down control of spinal nociception during distraction from pain, via the ACC-PAG-RVM-dorsal horn pathway. This indicates that the ACC signals high cognitive load associated with the task to the PAG, that recruits spinally-projecting cells in the RVM. Analgesia could be achieved through disinhibition of spinally-projecting OFF-cells (*21, 53, 54*), that inhibit dorsal horn neurons both directly via GABAergic and opioidergic projections to the primary afferents (*55, 56*) and also indirectly via local inhibitory interneuron pools at a spinal level (*46*) reflected in reduced BOLD signal in the *Spinal_noci_* cluster and activation of the *Spinal_int_* pool.

Previous human imaging studies have provided evidence for a role of the locus coeruleus in attentional analgesia (*16, 18*). We replicate some of those findings in showing activity in the LC related to both task and thermal stimulus as well as interactions between the LC and RVM that were modulated by the interaction between task and temperature. However, we neither found evidence for an interaction between task and temperature nor for a correlation with analgesic effect in the LC that we reported in our previous studies (*16, 18*). We also could not demonstrate altered connectivity between the LC and the spinal cord during the paradigm as we anticipated given its known role in descending pain modulation (*18, 22, 26, 28, 44, 57*). It is likely that the brainstem focussed slice prescription used previously is necessary for capturing sufficient signal from the LC, and that extending slice coverage compromised signal fidelity in these small brainstem nuclei. The noradrenergic manipulation with reboxetine did show a significant analgesic effect which was independent of task difficulty. This indicates that this dose of reboxetine is capable of altering baseline gain in the nociceptive system, but has no selective effect on attentional pain modulation. Reboxetine also modulated a task-dependent connection between ACC and PAG, though this did not appear to influence task performance and so its behavioural significance is uncertain. In interpreting these findings one potential explanation is that noradrenaline is not involved in attentional analgesia, however it could also be because of a ceiling effect where the reuptake inhibitor cannot increase the noradrenaline level any further during the attentional task. In this sense an antagonist experiment, similar to that was used to examine the role of the opioids, would be ideal. However, selective alpha2-antagonists are not used clinically and even experimental agents like Yohimbine have a number of issues that would have confounded this study in that they cause anxiety, excitation and hypertension. Therefore, we conclude that were not able to provide any additional causal evidence to support a role of the LC in attentional analgesia, but this likely reflects a limitation of our approach and lack of good pharmacological tools to resolve the influence of this challenging target.

This combination of simultaneous whole CNS imaging with concurrent thermal stimulation and attentional task in the context of pharmacological manipulation, has enabled the definition of long-range network influences on spinal nociceptive processes and their neurochemistry. An important aspect of this approach is that it has enabled the linkage between a large body of fundamental pain neuroscience that focussed on primary afferent to spinal communication and brainstem interactions (nociception) which can be directly integrated to the findings of whole CNS human imaging. This also offers novel opportunities for translational studies to investigate mechanisms and demonstrate drug target engagement. The finding that it is the effective connectivity of these networks that is of importance in the mediation of the effect of attention and the influence of the opioid antagonist reflects recent observations from large scale studies relating psychological measures to functional connectivity (e.g. (*58*)). In patient populations this focus on long range connectivity may help to differentiate between processes leading to augmented nociception and/or altered perception and control (e.g. in fibromyalgia (*34*)). Finally, we note that the location of the observed interaction between task and temperature indicates that cognitive tasks are integrated to act at the earliest level in the nociceptive transmission pathway introducing the novel concept of spinal psychology.

## Methods

### DATA ACQUISITION

#### Participants

Healthy volunteers were recruited through email and poster advertisement in the University of Bristol and were screened via self-report for their eligibility to participate. Exclusion criteria included any psychiatric disorder (including anxiety/depression), diagnosed chronic pain condition (e.g. fibromyalgia), left handedness, recent use of psychoactive compounds (e.g. recreational drugs or antidepressants) and standard MRI-safety exclusion criteria.

The study was approved by the University of Bristol Faculty of Science Human Research Ethics Committee (reference 23111759828). An initial power analysis was done to determine the sample size using the fmripower software (*59*). Using data from our previous study of attentional analgesia ((*16*), main effect of task contrast in the periaqueductal grey matter mask) we designed the study to have an 80% power to detect an effect size of 0.425 (one sample t-test) in the PAG with an alpha of 0.05 requiring a cohort of 40 subjects. Of fifty-seven subjects screened, two were excluded for claustrophobia, three were excluded for regular or recent drug use (including recreational), and five were excluded due to intolerance of the thermal stimulus. This was defined as high pain score (≥ 8/10) for a temperature that should be non-nociceptive (<43 °C). In addition, six participants withdrew from the study as they were unable to attend for the full three visits. One participant had an adverse reaction (nausea) to a study drug (naltrexone) and dropped out of the study. One subject was excluded for being unable to perform the task correctly. Thirty-nine participants completed all three study visits (mean age 23.7, range [18 - 45] years, 18 females).

#### Calibration of temperature and task velocity

In the first screening/calibration visit, the participants were briefed on the experiment and gave written informed consent. The participants were familiarised with thermal stimulation by undergoing a modified version of quantitative sensory testing (QST) based on the DFNS protocol (*60*). QST was performed using a Pathway device (MEDOC, Haifa, Israel) with a contact ATS thermode of surface area 9cm^2^ placed on the subject’s left forearm (corresponding to the C6 dermatome). Subsequently, the CHEPS thermode (surface area 5.73cm^2^) was used at the same site to deliver a 30 second hot stimulus, to determine the temperature to be used in the experimental visits. Each stimulus consisted of a plateau temperature of 36 to 45°C, and approximately thirty pseudorandomised “heat spikes” of 2, 3, or 4 degrees superimposed on the plateau, each lasting less than a second. Participant were asked to rate the sensation they felt during the whole stimulation period, on a scale from 0 (no pain) to 10 (the worst pain imaginable). The temperature which consistently produced a pain rating of 6 out of 10 was used for the noxious stimulation in the experiment. If the participant only gave pain scores lower than 6, then the maximum programmable plateau temperature of 45°C was used, but with higher temperature spikes of 3, 4 and 5 degrees above, reaching the highest temperature allowed for safety (50°C maximum).

The session also included a calibration of the rapid serial visual presentation (RSVP) task (*29*), where participants were asked to spot the number 5 among distractor characters. The task was repeated 16 times at different velocities (i.e. different inter-character intervals) in pseudorandom order, ranging from 32 to 256ms. To identify the optimal speed for the hard version of the RSVP task (defined as 70% of each subject’s maximum d’ score), the d’ scores for the different velocities were plotted and the curve fit to a sigmoidal function, using a non-linear least squares fitting routine in Excel (Solver). Once parameterised, the target speed for 70% performance was recorded for subsequent use during the imaging session.

#### Imaging sessions

Following the screening/calibration session, participants returned for three imaging sessions, spaced at least a week apart. Participants underwent drug screening (questionnaire) and pregnancy testing. After eating a light snack, they were given either an inert placebo capsule, naltrexone (50mg) or reboxetine (4mg) according to a randomised schedule. The tablets were encased in identical gelatine capsules and dispensed in numbered bottles prepared by the hospital pharmacy (Bristol Royal Infirmary, University Hospitals Bristol and Weston NHS Foundation Trust).

One hour after drug dosing, calibration of the RSVP task was repeated (to control for any effect on performance). Before scanning, participants received the high thermal stimulus at their pre-determined temperature, which they rated verbally. If the rating was 6±1, the temperature was kept the same, otherwise it was adjusted accordingly (up or down). Neither reboxetine nor naltrexone caused a significant change in pain perception or task velocity during the calibration, as verified with paired t tests (placebo versus reboxetine and placebo versus naltrexone, see **Supplementary Figure**). On average, the plateau temperature used for high temperature stimuli was 43.8± 1.25°C. The median inter-stimulus interval for the hard RSVP task was 48ms, range [32-96].

In the MRI scanner, participants performed the RSVP task at either difficulty level (easy or hard) whilst innocuous (low) or noxious (high) thermal stimuli were delivered concurrently to their left forearm. The four experimental conditions (*easy|high, hard|high, easy|low, hard|low*), were repeated four times each, in a pseudo-random order. The hard version (70% d’ performance) of the task and the high (noxious) thermal stimulus were calibrated as described above. In the easy version of the task the inter-character presentation speed was always set at 192ms, except when a participant’s hard task velocity of was equal or slower than 96ms, whereby the easy task was set to 256ms. The low (innocuous) thermal stimulus was always set to be a plateau of 36 °C with spikes of 2, 3 and 4°C above this baseline. Participants performed the task (identifying targets) and provided pain ratings 10 seconds after the end of each experimental block on a visual analogue scale (0-100), using a button box (Lumina) held in their dominant (right) hand.

#### Acquisition of functional images

Functional images were obtained with a 3T Siemens Skyra MRI scanner, and 64 channel receive-only head and neck coil. After acquisition of localiser images, a sagittal volumetric T1-weighted structural image of brain, brainstem and spinal cord was acquired using the MPRAGE pulse sequence, (TR = 2000ms, TE = 3.72ms, TI = 1000 ms, flip angle 9°, field of view (FoV) 320 mm, GRAPPA acceleration factor = 2) and 1.0mm isotropic resolution. Blood oxygenation level dependent (BOLD) functional data was acquired axially from the top of the brain to the intervertebral disc between the C6 and C7 vertebral bodies, with TR = 3000ms, TE = 39ms, GRAPPA acceleration factor = 2, flip angle 90°, FoV 170 mm, phase encoding direction P>>A, matrix size 96 by 96. Slices were positioned perpendicular to the long axis of the cord for the C5-C6 spinal segments, whilst still maintaining whole brain coverage, and had an in-plane resolution of 1.77 × 1.77 mm and slice thickness of 4mm and a 40% gap between slices (increased to 45-50% in taller participants). To determine the optimal shim offset for each slice, calibration scans were acquired cycling through 15 shim offsets. For the caudal 20 slices covering from spinal cord to medulla, manual inspection of images determined the optimal shim offset to be used for each subject (*31*). The remaining supraspinal slices were acquired with the first and higher order shim offsets determined using the scanner’s automated routine. During scanning, cardiac and respiratory processes were recorded using a finger pulse oximeter (Nonin 7500) and pneumatic respiratory bellows (Lafayette), respectively. These physiological signals and scanner triggers were recorded using an MP150 data acquisition unit (BIOPAC, Goleta, CA), and converted to text files for subsequent use during signal modelling.

### DATA ANALYSIS

#### Analysis of pain scores

Pain scores recorded during the experiment were investigated collectively for the three visits using a three-way ANOVA in Prism version 8 for Windows (GraphPad Software, La Jolla, California). Any significant interaction was further investigated with two separate three-way ANOVAs (placebo versus naltrexone and placebo versus reboxetine). Finally, each drug condition was analysed individually with three separate two-way ANOVAs. Two-tailed post-hoc tests were used to further investigate any interactions.

#### Pre-processing of functional data and single-subject analysis

Functional data were divided into spinal cord and brain/brainstem, by splitting at the top of the odontoid process (dens) of the 2^nd^ cervical vertebra. The resulting two sets of images underwent separate, optimised, pre-processing pipelines.

Spinal cord data was motion corrected with AFNI 2dImReg (*61*), registering all time points to the temporal mean. Data was smoothed with an in-plane 2D Gaussian smoothing kernel of 2mm × 2mm FWHM, using an in-house generated script. The Spinal Cord Toolbox (SCT, v4.1.1) was then used to create a 25mm diameter cylindrical mask around the entire cord to crop the functional data. The SCT was also used to segment the cord from the cerebrospinal fluid (CSF) and register functional images to the PAM50 template (*62*). Manual intervention was necessary to ensure accurate cord segmentation. The inverse warping fields generated by the registration of spinal cord fMRI data to the PAM50 template were used to warp a PAM50 CSF mask to subject space. The mask was then used to create a CSF regressor for use during correction for physiological noise during first level FEAT analysis (part of FSL, (*63*))

Brain functional data was pre-processed and analysed in FEAT. Pre-processing included smoothing with a 6mm Gaussian kernel, and motion correction with MCFLIRT (*64*). Functional data was unwarped with a fieldmap using FUGUE (*65*), then co-registered to the subject’s structural (T1-weightedscan using boundary-based registration (*66*). Structural scans were registered to the 2mm MNI template using a combination of linear (FLIRT, (*67*)) and non-linear (FNIRT, (*68*)) registration with 5mm warp resolution.

Physiological noise correction was conducted for the brain and spinal cord (*69, 70*) within FEAT. Cardiac and respiratory phases were determined using a physiological noise model (PNM, part of FSL), and slice specific regressors determined for the entire CNS coverage. Subsequently these regressors (which are 4D images) were split at the level of the odontoid process, to be used separately for brain and spinal cord physiological noise correction. For the brain data the PNM consisted of 32 regressors, with the addition of a CSF regressor for the spinal cord, giving a total of 33 regressors for this region.

All functional images were analysed using a general linear model (GLM) in FEAT with high-pass temporal filtering (cut-off 90s) and pre-whitening using FILM (*71*). The model included a regressor for each of the experimental conditions (*easy|high*, *hard|high*, *easy|low*, *hard|low*), plus regressors of no interest (task instructions, rating periods), and their temporal derivatives. Motion parameters and physiological regressors were also included in the model to help explain signal variation due to bulk movement and physiological noise. The experimental regressors of interest were used to build the following planned statistical contrasts: positive and negative main effect of temperature (high temperature conditions versus low temperature conditions and vice versa), positive and negative main effect of task (hard task conditions versus easy task conditions and vice versa), and positive and negative interactions.

#### Group analysis

We used a conservative approach to investigate differences in CNS activity in main effects and interactions due to administration of reboxetine or naltrexone. An initial analysis examined the brain, brainstem, and spinal cord activation in the planned contrasts (main effects of temperature, task, and their interaction) across all visits i.e. a “pooled” analysis. Individual subjects’ data were averaged using a within-subject “group” model (treating variance between sessions as a random effect), and resultant outputs averaged (across subjects) using a mixed effects model. This allowed the generation of functional masks, to use for investigation of differences between drug conditions.

Generalised psychophysiological interaction (gPPI) analysis (*72*) was used to assess effective connectivity changes between brain, brainstem, and spinal cord during the attentional analgesia experiment. The list of regions to be investigated were specified *a priori* on the basis of our previous study (*18*), and included the ACC, PAG, LC and RVM – to which was added the left side of the spinal cord at the C5/C6 vertebral level. Following partial unblinding to drug, an initial analysis was performed for the placebo visit, with the purpose of building functional localizers. Any significant differences in connectivity identified in the placebo arm, across the experimental conditions, were thus examined for effects of pharmacological interventions (i.e. drugs causing significant connectivity changes).

All first-level analyses, single group averages and pooled analyses were performed with the experimenter blind to the study visit (i.e. drug session). Subsequently, the experimenter was partially unblinded to the placebo visit to determine a functional localiser for the purpose of connectivity analysis, and to perform paired t tests between conditions. The experimenter was finally unblinded to all the visits for interpretation of results.

#### Pooled analysis – spinal cord

For each subject, parameter maps estimated for each contrast and each visit (i.e. drug session), were registered to the PAM50 template with SCT. Each contrast was then averaged across visits using a within-subject ordinary least squares (OLS) model using FLAME (part of FSL) from command line. The resulting average contrasts (registered to the PAM50 template) were each concatenated across subjects (i.e. each contrast had 39 samples). These were then investigated with a one-sample t-test in RANDOMISE, using a left C5-6 vertebral mask, derived from SCT. Results are reported with threshold free cluster enhancement (TFCE) P < 0.05 corrected for multiple comparisons, after a two-tailed test. Significant regions of activation from this pooled analysis were used to generate masks for subsequent comparison between conditions, using paired t-tests.

#### Pooled analysis – brainstem

Similar to the spinal cord, for each subject, parameter maps from the brainstem for each planned contrast and visit were averaged with an OLS model in FEAT software. The resulting average was the input to a between-subjects, mixed effects, one-sample t-test in FEAT. Subsequently, group activations for each of the six contrasts were investigated with permutation testing in RANDOMISE, using a probabilistic mask of the brainstem taken from the Harvard-Oxford subcortical atlas (threshold set to P=0.5). Results are reported with TFCE correction and P < 0.05, two tailed. Significant regions of activity were binarized and used as a functional mask for the between conditions comparison.

#### Pooled analysis – brain

Brain data was averaged and analysed with the same FEAT analyses that were applied in the brainstem. Following within subject averaging, group activity was assessed with a mixed effects two-tailed one sample t-test at the whole-brain level, with results reported for cluster forming threshold of Z > 3.1, and corrected cluster significance of P < 0.05. This produced maps of activity (one per planned contrast) that were then binarized to produce masks that were used in follow up paired t-tests.

#### Within subject comparison – paired tests

Paired t tests were performed to resolve potential changes in activity in reboxetine versus placebo and naltrexone versus placebo, separately. Design and contrast files for input in RANDOMISE were built in FEAT. A group file with appropriately defined exchangeability blocks was additionally defined. Permutation testing in RANDOMISE was used to assess group level differences between placebo and the two drugs, separately for brain, brainstem, and spinal cord. The investigation was restricted to the functional masks derived from the main effect analysis for each contrast.

#### Effective connectivity analysis (gPPI)

For connectivity analysis, functional data for brain, brainstem and spinal cord were pre-processed as previously described (*18*). Time-series for the seed region were extracted from the voxel of greatest significance identified in the analysis of the placebo session within the prespecified anatomical regions. In particular, data was extracted from the peak voxel responding to the main effect of temperature in the RVM and spinal cord, the main effect of task in the ACC, PAG and LC, and the task * temperature interaction in the spinal cord. The time-series were included in a GLM that also included the same regressors present in the first level main effects analysis i.e. regressors for the experimental conditions and all nuisance regressors (rating period, instruction, PNM, movement parameters). Interaction regressors were then built by multiplying the time-series by each of the experimental regressors, and the planned contrasts constructed (e.g. positive main effect of task). Apart from systematically varying the input physiological timeseries corresponding to different seed regions, models used for estimating connectivity for brain and spinal cord seeds were otherwise identical. Estimates of effective connectivity for the group were obtained with permutation testing with RANDOMISE, using as targets the ROI masks used for time-series extraction. For example, a gPPI analysis with an RVM seed timeseries, used PAG, LC, ACC, and a left C5-6 vertebral mask to estimate connectivity changes between brain/brainstem and spinal cord. Two-tailed paired t-tests were used to detect differences (TFCE, P<0.05) between drug visits in the significant connections, as described above.

## Authors Contributions

Conceptualization: VO, RM, AEP, JCWB.

Methodology: RHD, JCWB

Investigation: VO, AEP, JCWB

Visualization: VO, AEP, JCWB

Supervision: JCWB, AEP, RM

Writing – original draft: VO, AEP, JCWB

Writing – review & editing: VO, RM, AEP, JCWB

## Acknowledgements

The authors would like to thank Aileen Wilson (Lead Research Radiographer, CRiCBristol) for her support in running experiments, and the subjects who kindly agreed to take part. This research was funded in whole, or in part, by the Wellcome Trust [203963/Z/16/Z; and 088373/Z/09/A] and Medical Research Council [MR/N026969/1]. For the purpose of Open Access, the author has applied a CC BY public copyright licence to any Author Accepted Manuscript version arising from this submission.

---

**Supplementary Figure 1.**
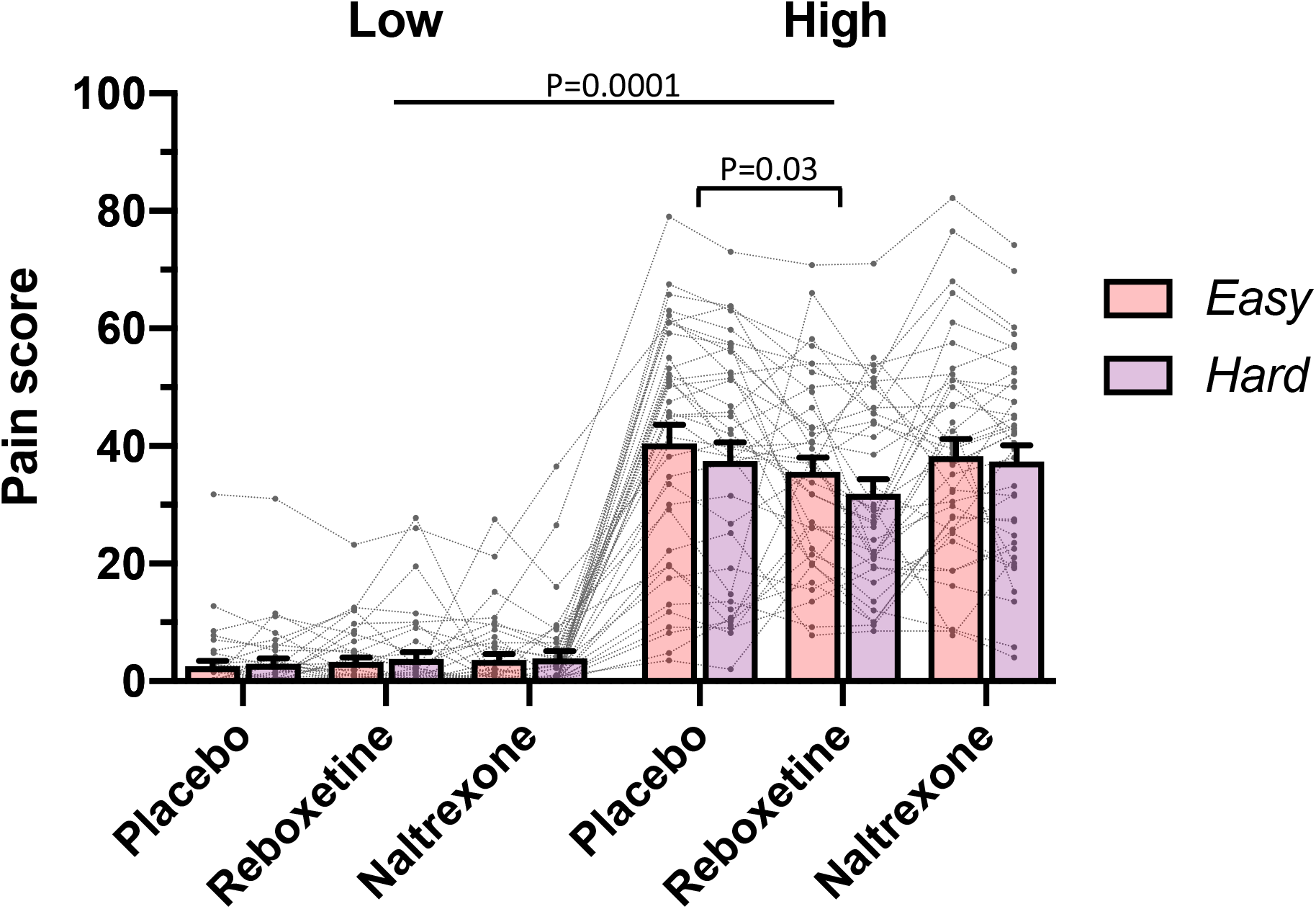
Pain scores under the four experimental conditions (i.e. easy|low, hard|low, easy|high and hard|high), across the three drugs for each of the 39 subjects. A first level, three-way mixed effects ANOVA revealed the expected main effect of temperature (F (1,38) = 221, P=0.0001), main effect of task (F (1,38) = 4.9, P=0.03) and importantly a task*temperature interaction (F (1, 38) = 10.5, P = 0.0025). The first level analysis also showed a drug*temperature interaction on pain ratings (F (2, 76) = 3.2, P = 0.04). To further investigate the drug*temperature interaction, two second level three-way mixed effects ANOVAs were conducted for placebo vs reboxetine and placebo vs naltrexone. For reboxetine versus placebo, a drug*temperature interaction was revealed (F (1, 38) = 5.060, P = 0.03), with lower pain scores in high temperature condition in the reboxetine arm, indicating an analgesic effect of the drug. No interactions were observed in the ANOVA contrasting naltrexone with placebo. Mean+SEM with individual participants data.

**Supplementary Figure 2.**
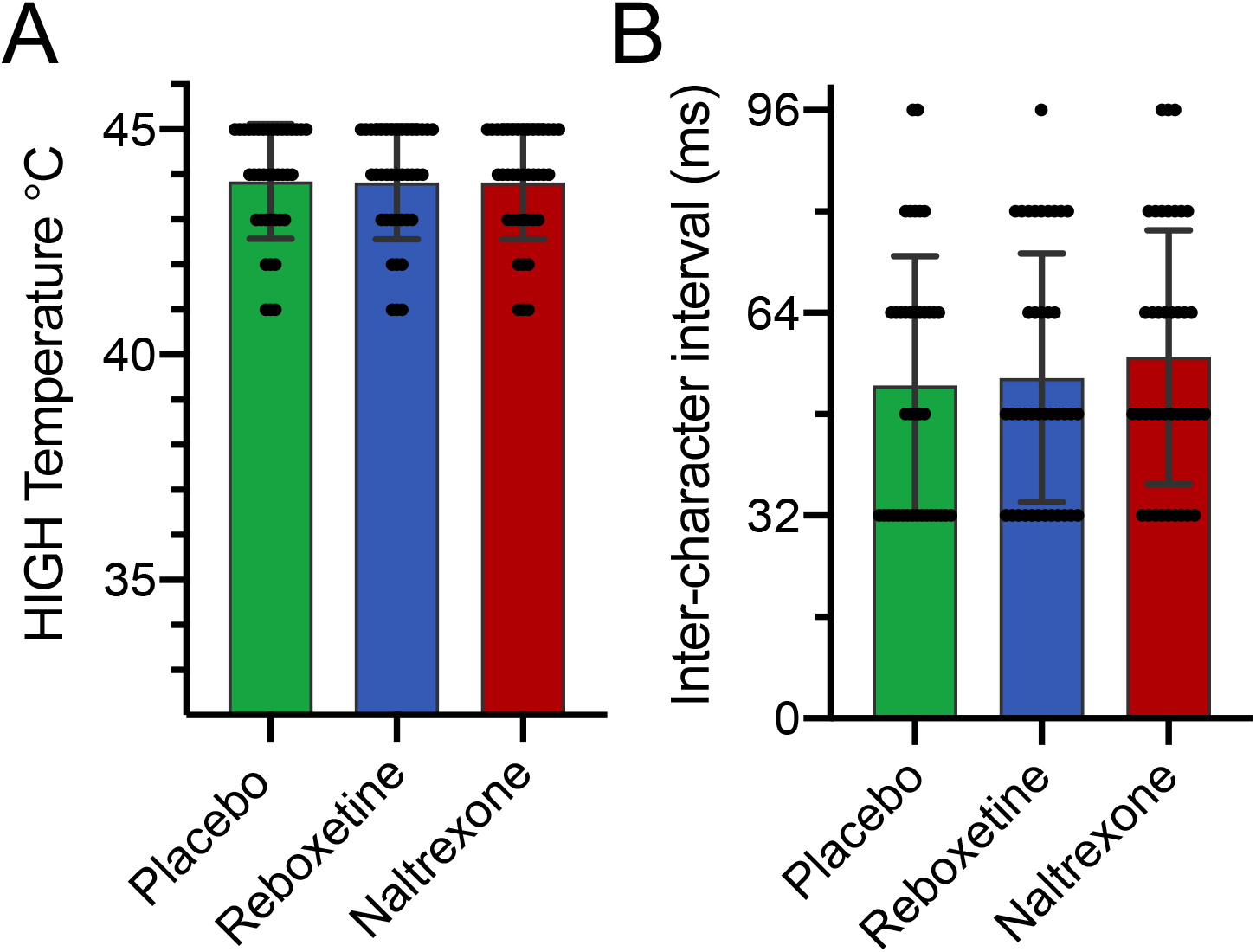
Temperature delivered and task speed across the three drug conditions. (A) Administration of Reboxetine or Naltrexone did not change the individually calibrated HIGH thermal stimulus required to evoke a 6/10 pain score (Mean ± SD). (B) Similarly, drug administration had no effect on RSVP task speed as reflected in the inter-character presentation interval. (Mean ±SD, Friedman tests NS).

**Supplementary Figure 3.**
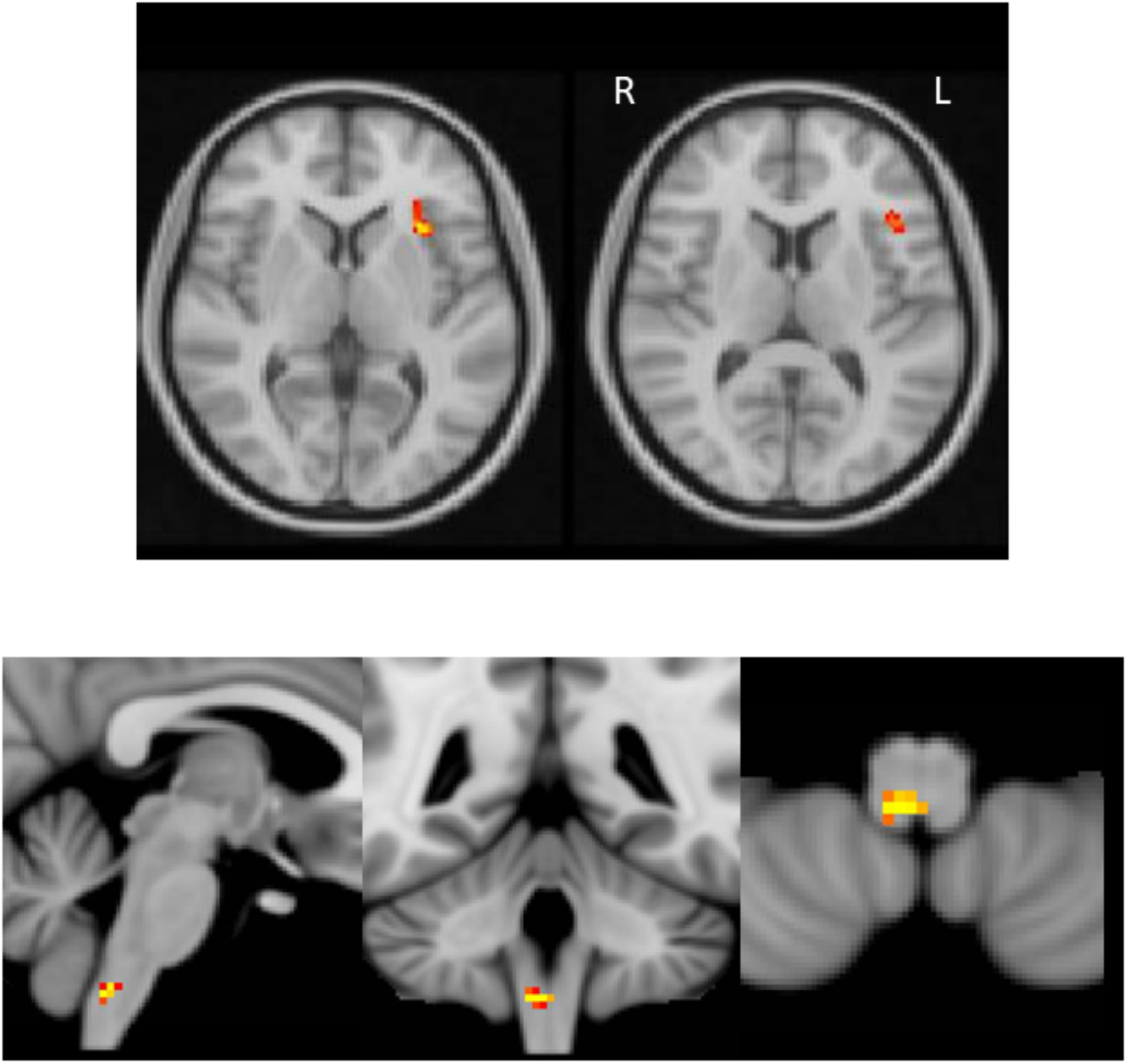
Anterior Insula and medulla response after Naltrexone administration. (A) The anterior insula responded more strongly in the naltrexone than in the placebo in the main effect of task (obtained with permutation testing with a main effect of task mask, obtained from the pooled analysis). (B) A cluster in the lower medulla responded more strongly in the naltrexone than in the placebo main effect of temperature. Result obtained with permutation testing (using a main effect of temperature brainstem mask, obtained from the pooled analysis).

## Supplementary Table

**Supplementary Table 1.**
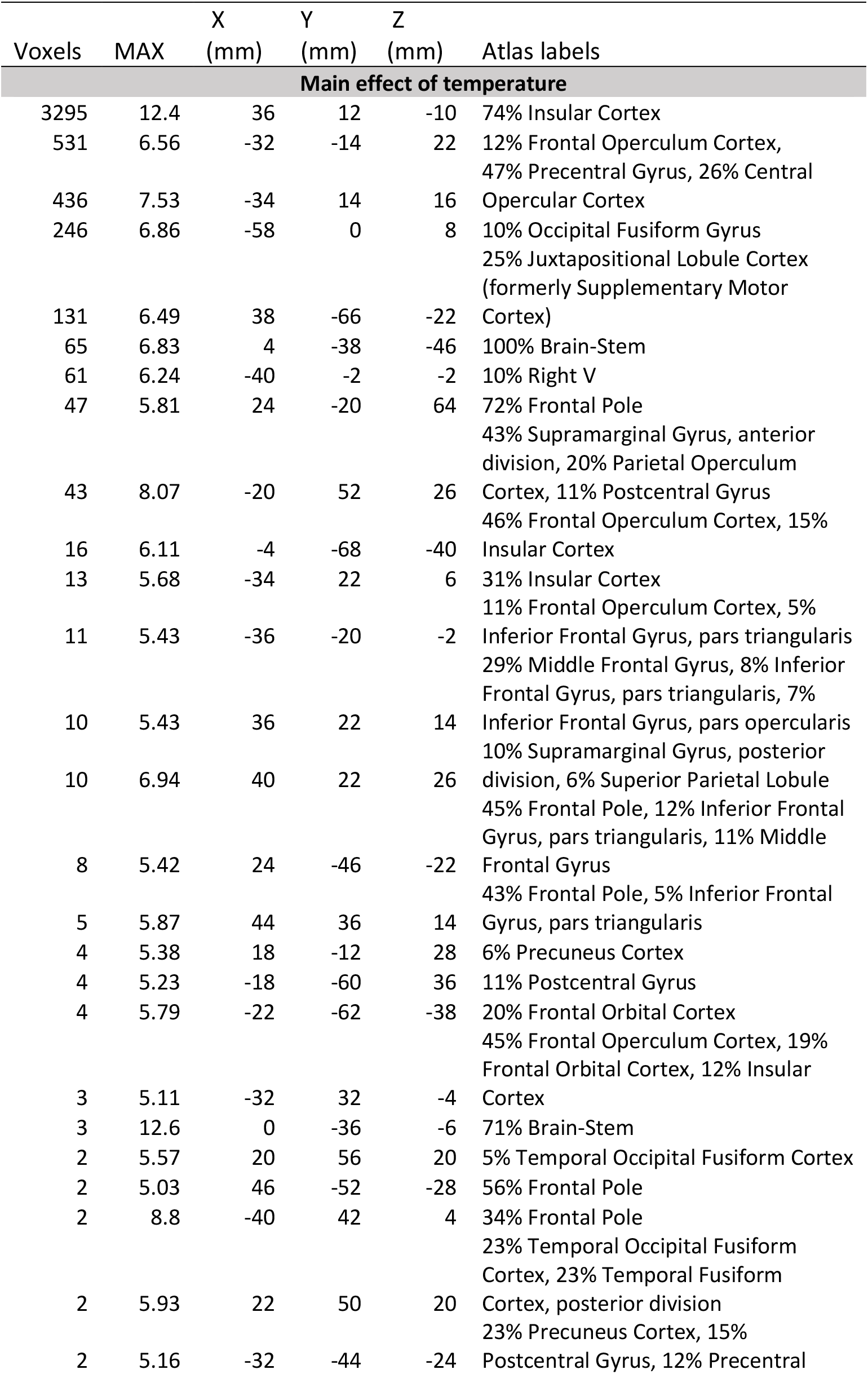
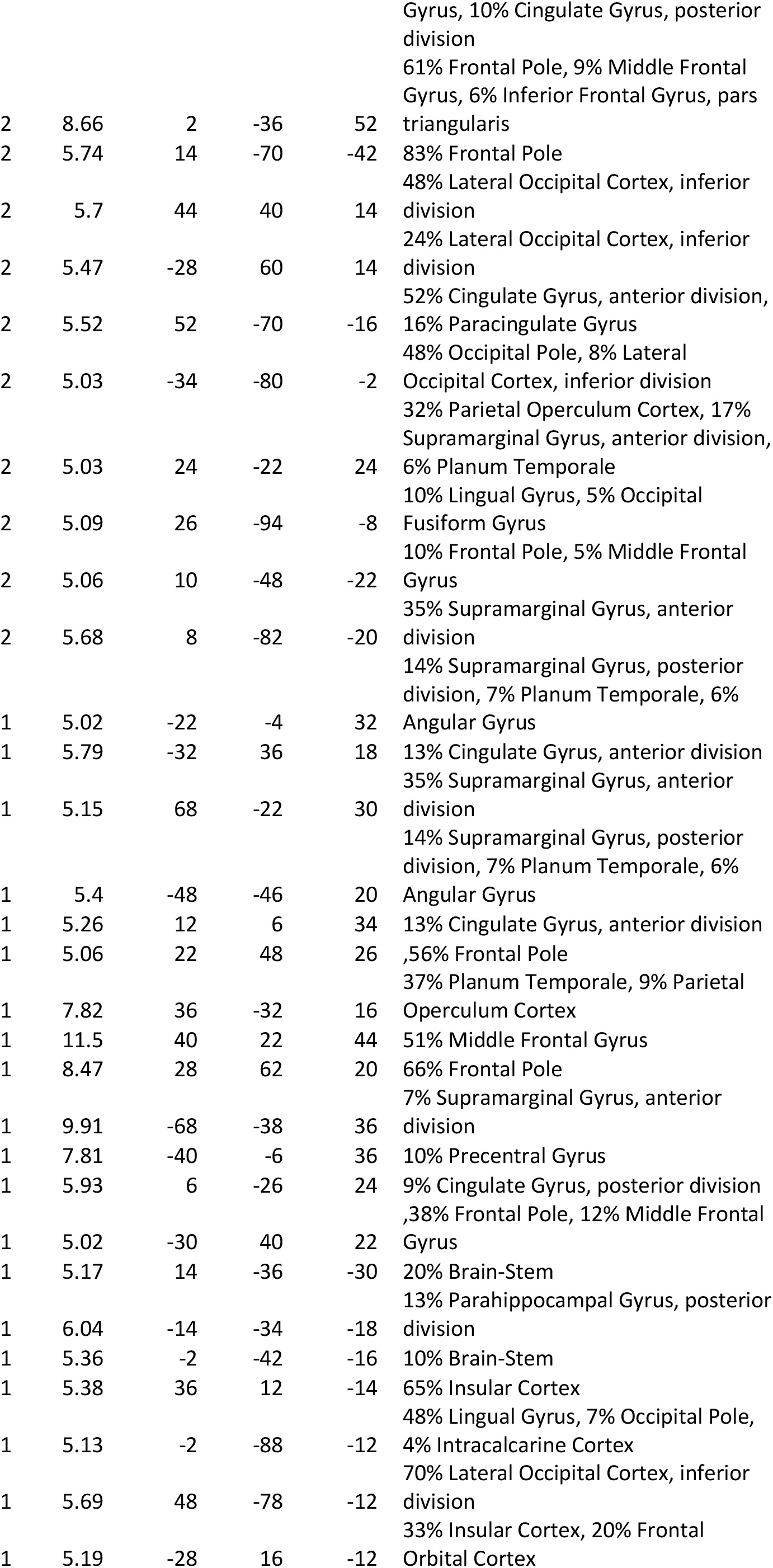
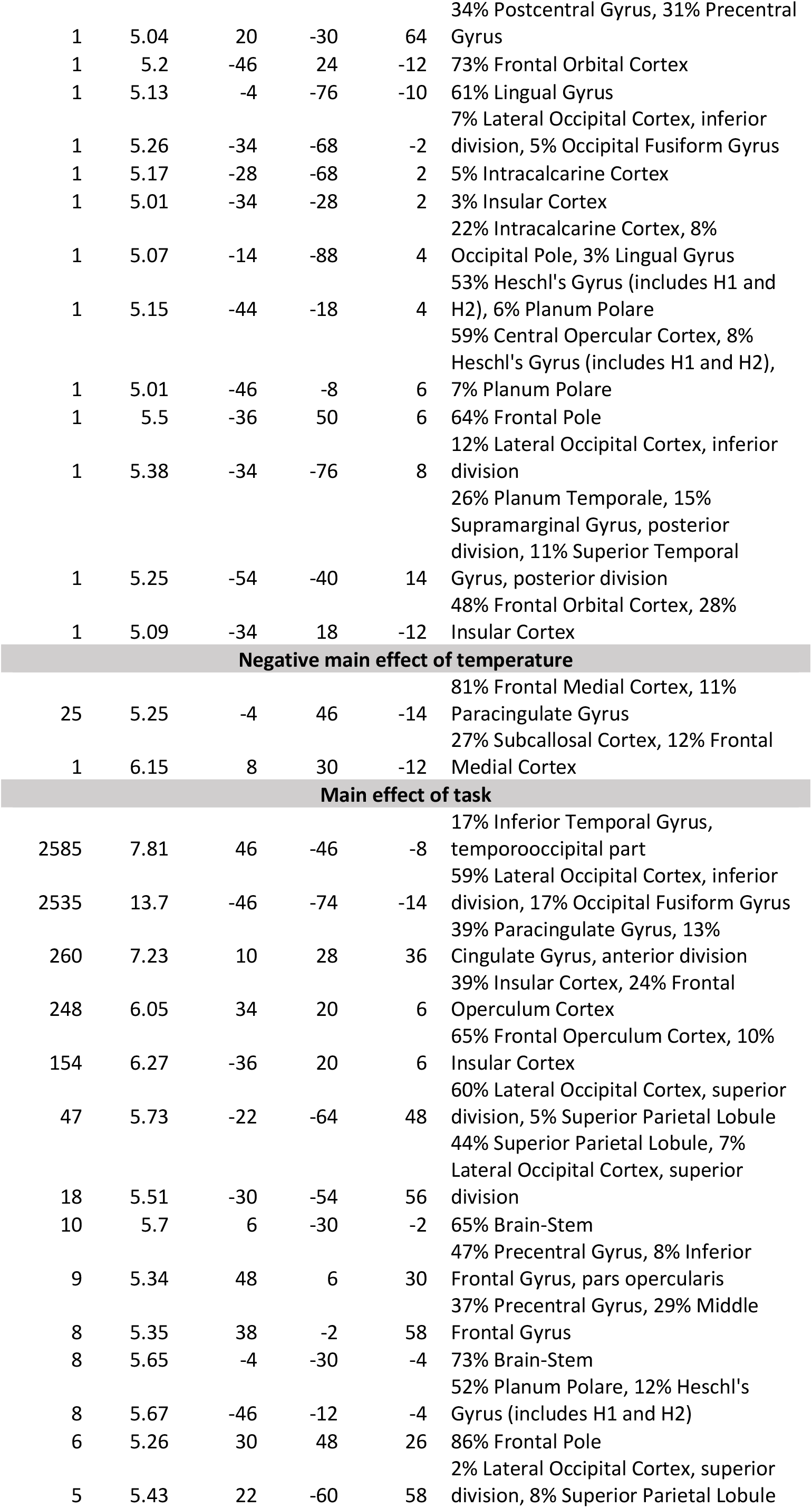
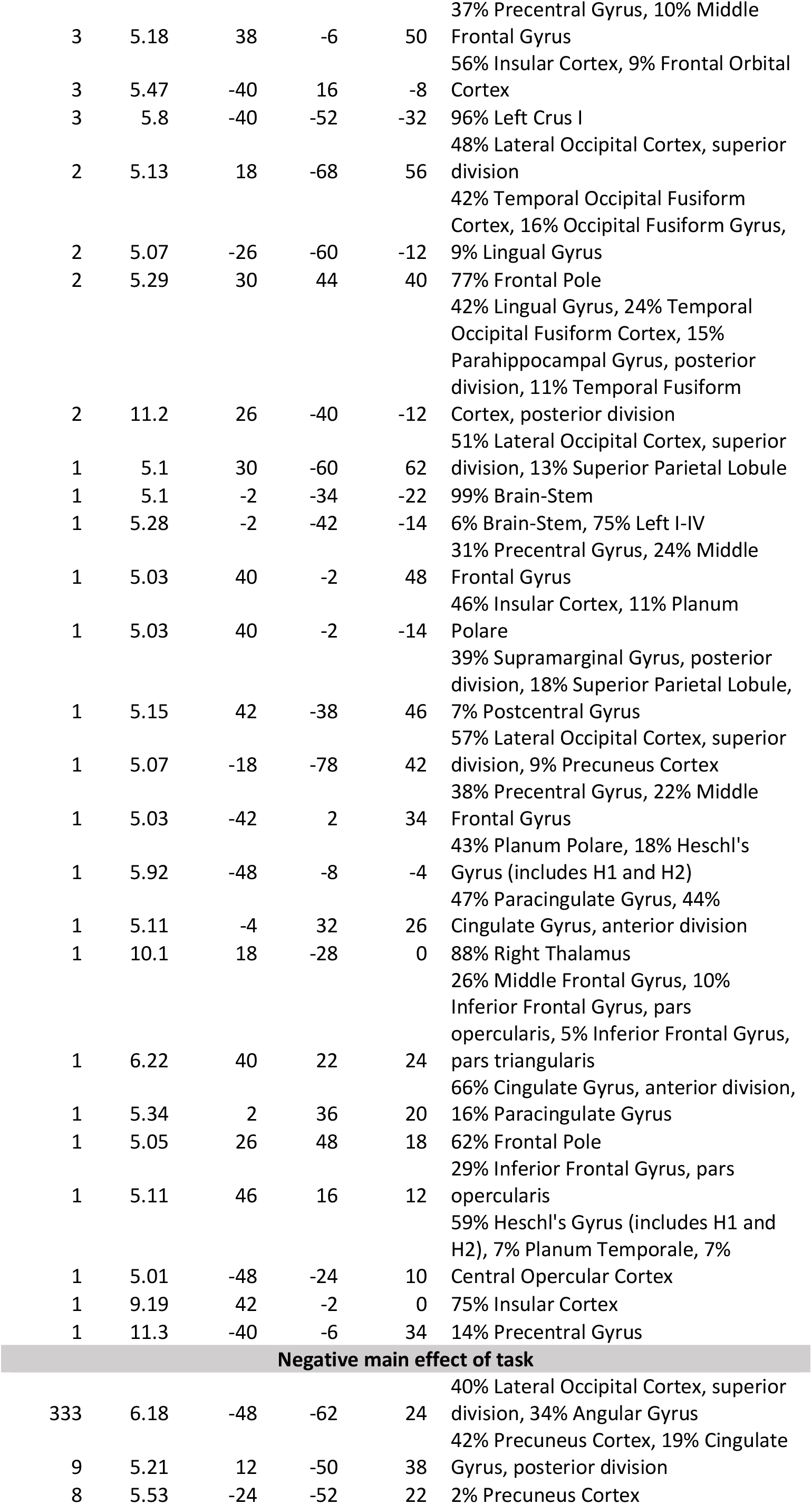
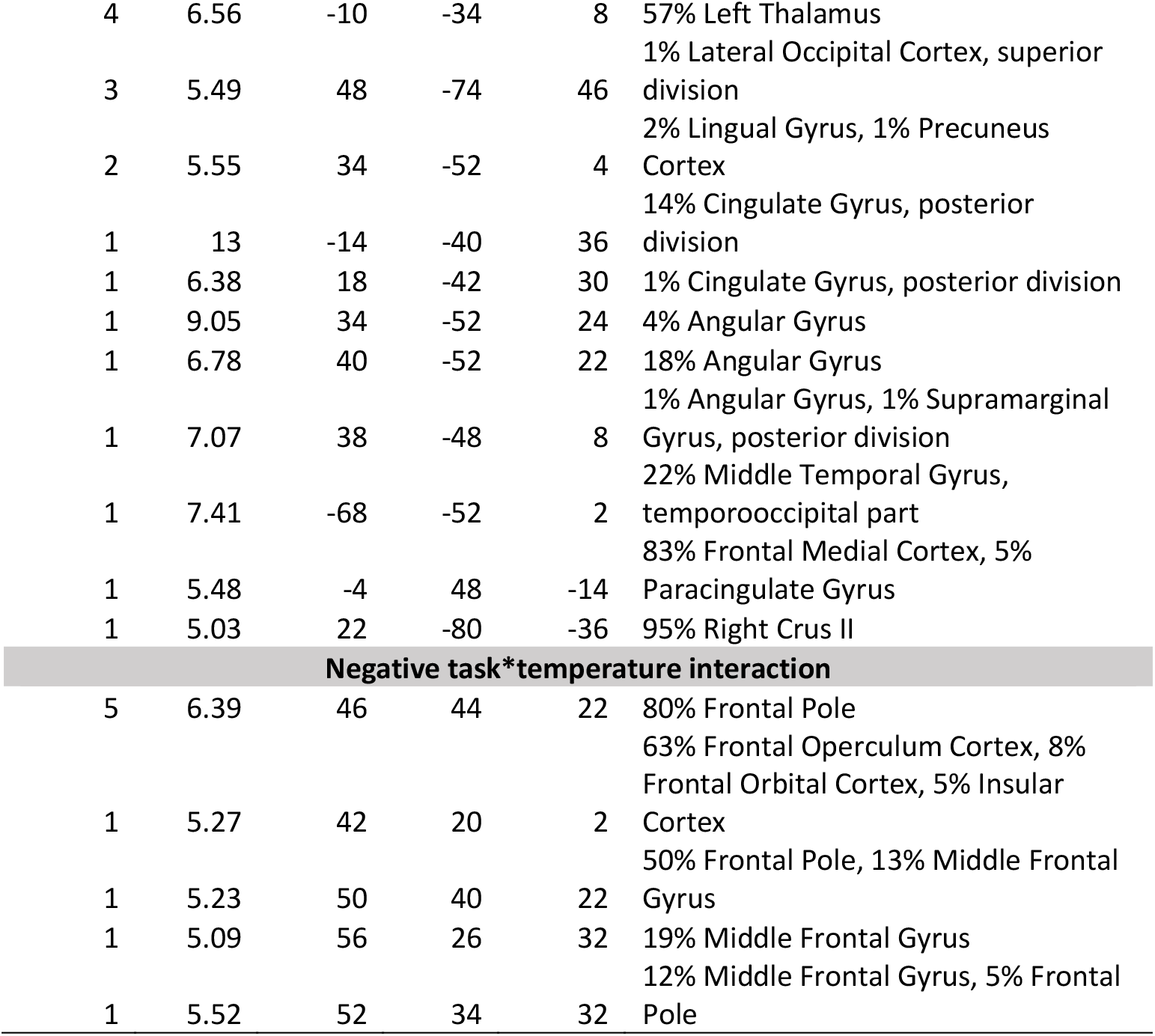
Results from main effect analyses in the whole brain, across the three drug conditions. Obtained with cluster-forming threshold Z>3.09 and cluster-corrected p<0.05. The tables were created with Autoaq (part of FSL), with atlas labels based on the degree of overlap with probabilistic atlases (Harvard Oxford Cortical Structural Atlas, Harvard Oxford Subcortical Structural Atlas, Cerebellar Atlas in MNI152 space after normalization with FNIRT).

